# Differences of endogenous polyamines and putative genes associated with paraquat resistance in goosegrass (*Eleusine indica L.*)

**DOI:** 10.1101/618074

**Authors:** Qiyu Luo, Jiping Wei, Zhaoxia Dong, Xuefeng Shen, Yong Chen

**Affiliations:** Department of Crop Cultivation and Farming System, South China Agricultural University, Guangzhou, Guangdong, China

## Abstract

**Background:** Paraquat is one of the most effective herbicides used to control weeds in agricultural management, while the pernicious weed goosegrass (*Eleusine indica*) has evolved resistance to herbicides, including paraquat. Polyamines provide high-level paraquat resistance in many plants. In the present study, we selected three polyamines, namely, putrescine, spermidine, and spermine, as putative genes to investigate their correlation with paraquat resistance by using paraquat-resistant (R) and paraquat-susceptible (S) goosegrass populations.

**Results:** There was no significant difference in the putrescine nor spermine content between the R and S biotypes. However, 30 and 90 min after paraquat treatment, the spermidine concentration was 346.14-fold and 421.04-fold (P < 0.001) higher in the R biotype than in the S biotype, but the spermidine concentration was drastically reduced to a marginal level after 90 min. Since the transcript level of *PqE* was low while the spermidine concentration showed a transient increase, the *PqE* gene was likely involved in the synthesis of the paraquat resistance mechanism, regulation of polyamine content, and synthesis of spermidine and spermine. *PqTS1*, *PqTS2,* and *PqTS3* encode transporter proteins involved in the regulation of paraquat concentration but showed different transcription patterns with synchronous changes in polyamine content.

**Conclusion:** Endogenous polyamines (especially spermidine) play a vital role in paraquat resistance in goosegrass. *PqE, PqTS1, PqTS2*, and *PqTS3* were speculated on the relationship between polyamine metabolism and paraquat resistance. To validate the roles of *PqE, PqTS1, PqTS2*, and *PqTS3* in polyamine transport systems, further research is needed.

## Introduction

Paraquat is a haloid salt [1] that is widely used as a non-selective herbicide to control weeds throughout the world [2-4]. Paraquat resistance in plant cells is determined by the uptake, efflux, sequestration, detoxification, and catabolism of the reactive oxygen species (ROS) generated by paraquat.^3^ ROS can cause tissue damage by reacting with lipids in cellular membranes and nucleotides in DNA [5]. Two hypotheses have been proposed for paraquat resistance in plants: One is an increase in the capability of plant cells to scavenge ROS, while the other is the ability of plant cells to sequester paraquat away from target sites in their chloroplasts [6]. Although there are numerous data available on paraquat resistance [7], the molecular mechanism has yet to be elucidated [2]. Based on the scant data available, paraquat resistance in model plants involves a single major gene mechanism, but the molecular detail of this mechanism is not fully understood [8].To the best of our knowledge, there are no studies on the molecular mechanism of paraquat resistance in weeds.

Polyamines, including putrescine, spermidine and spermine, are small aliphatic polycations that are classified as growth regulators, although the specific mechanisms have not been well elucidated [9]. Exogenous application of polyamines provides high-level resistance against paraquat toxicity in a range of plant organs [4], such as the roots of intact maize seedlings [10], leaf discs of sunflower [11], and leaves of rice [12]. Endogenous polyamine probably plays a vital role in paraquat resistance in plants. However, there is little information available on the mechanism of endogenous polyamine activities in paraquat resistance.

Several genes encoding putative components of polyamine transport systems have been identified in bacteria. Quantification of the polyamine contents in paraquat-resistant (R) and paraquat-susceptible (S) *Arctotheca calendula* revealed a possible role of polyamines and/or polyamine transporters in paraquat resistance [13]. The structure of paraquat is similar to that of polyamines, and both can use the same transport system in their hosts [14]. Since polyamines can be transported into the vacuoles, a polyamine transporter likely mediates this transport process [15]. A study on the role of transporters in the paraquat resistance of horseweed showed that the 12-kDa *E. coli* multidrug transporter (EmrE) and an amino acid transporter (CAT4) were related to paraquat resistance, and the CAT4 transporter had affinity for substances with a charge distribution similar to that of polyamine molecules [1]. Despite extensive studies on polyamine metabolism, the functions of polyamine-related genes in plant resistance to paraquat remain unclear.

Herbicides are the most widely used tool for weed management in large production areas where hand-weeding is neither efficient nor economical [16]. To date, nearly 30 species of R weed species have been reported worldwide [17]. Goosegrass (*Eleusine indica (L.)* Gaertn) is a pernicious weed species with a cosmopolitan distribution and is usually controlled by paraquat [18-19]. However, R biotypes of goosegrass have arisen due to the widespread use of the herbicide [20]. Our previous study identified 28 genes related to paraquat resistance in goosegrass. Ten of these genes were related to polyamines, and the remaining 18 genes were related to polyamine transporters [21]. Thus, further research on the role of polyamine transporter genes in paraquat resistance in goosegrass can be used to understand the molecular mechanism of paraquat resistance.

We focused on the genes *PqE, PqTS1, PqTS2,* and *PqTS3*, which were significantly upregulated in S and R biotypes 80 min after paraquat treatment, with high resistant character values (RQ/SQ) of +1.07, +1.01, +1.20, and +1.03, respectively [21]. Transcriptome analysis revealed *PqE* as the putative gene related to polyamine synthesis, while *PqTS1, PqTS2,* and *PqTS3* were identified as the putative genes associated with paraquat resistance in goosegrass. In this study, S and R biotypes of goosegrass were treated with paraquat for 180 min. We used real-time PCR (RT-PCR) to validate high-throughput sequencing data from S and R goosegrass, and the transcript levels of *PqE, PqTS1, PqTS2,* and *PqTS3* were determined after treatment. We also quantified endogenous polyamine contents in both biotypes using high-performance liquid chromatography (HPLC). Based on the results, we speculated on the relationship between the genes related to polyamine metabolism and paraquat resistance. These findings offer explanations of the molecular mechanisms of paraquat resistance.

## Materials and methods

### Plant materials and experimental treatments

The R biotype was collected from the Teaching and Research Farm (113°40′E, 22°80′N) in Panyu District, Guangzhou, China. Papaya (*Carica papaya* L.) and banana (*Musa nana* Lour.) are cultivated in this area, and paraquat has been used to control weeds continuously for 20 years. The S biotype was collected from the campus of South China Agricultural University (113°36′E, 23°16′N), where there is no history of paraquat use. The R biotype was confirmed prior to performing experiments, and the level of resistance in the R biotype was 59.48-fold higher than that in the S biotype [22]. For both biotypes, goosegrass seedling cultivation and paraquat treatment were performed as follows: seeds were scarified with sandpaper, sterilized for 10 min in 3% NaClO, washed three times, imbibed for 24 h in double-distilled water, and then germinated in plastic boxes (22×15.5×7 cm) containing a 2:1:1 mixture of soil:peat:sand in a growth chamber at 34°C/28°C (day/night) with a 12 h photoperiod and a light intensity of 800±200 µE m^-2^ s^-1^. Fourteen days after sowing (DAS), seedlings of both the S and R biotypes of goosegrass were transplanted into 24 pots (9×7 cm), each containing 6 plants. At 21 DAS, seedlings of both the S and R biotypes at the five leaf stage were sprayed with 0.6 kg·ai ha^-1^ paraquat (Syngenta Corporation, Shanghai, China) (the recommended rate) using a 3WP-2000 spray tower (Nanjing Research Institute for Agricultural Mechanization, Ministry of Agriculture, Nanjing, China). After 0, 30, 60, 90, 120, and 180 min, the aboveground parts of goosegrass were collected from treated S and R biotype seedlings sprayed with paraquat. These samples were collected with sterilized scissors and transferred into 10 ml centrifuge tubes using sterilized tweezers. All samples were flash frozen in liquid nitrogen and stored at −80°C pending analysis. Each treatment had 4 replicates.

### Real-time PCR

Total RNA was isolated using a EasyPure Plant RNA Kit (TransGen Biotech, Beijing,China). The concentration and purity of total RNAs were evaluated by ultraviolet (UV) light absorption spectra and the A260/A280 ratio (Appendix 1). Total RNA was used to synthesize cDNA with a PrimeScript II 1st Strand cDNA Synthesis Kit (D6210A; TaKaRa, Dalian, China) according to the manufacturer’s protocol. RT-PCRs were carried out using a Bio-Rad iCycler (CFX96, Bio-Rad, Santa Rosa, California, USA) with the following cycling conditions: 95°C for 30 s (20°C s^-1^) and 45 cycles of 95°C for 5 s (20°C s^-1^) and 60°C for 20 s (20°C s^-1^), followed by one cycle of 95°C for 0 s (20°C s^-1^), 60°C for 15 s (20°C s^-1^), and 95°C × 0 s (0.1°C s^-1^). Data were analysed with Bio-Rad CFX Manager Software 1.6 using the 2^-ΔΔCt^ (threshold cycle values) method. Relative PCR product levels were calculated from amplification and dissolution curves for each putative gene. The EST database was searched using the Basic Local Alignment Search Tool (BLAST) [23]. Multiple sequence alignments of amino acid sequences were conducted using CLUSTALW [24]. The high-throughput sequencing results for the 4 selected putative genes (*PqE, PqTS1, PqTS2,* and *PqTS3*) associated with paraquat resistance in goosegrass were obtained from a previous study [21]. The goosegrass *Actin* gene was amplified as an internal control. The gene-specific primers used for RT-PCR were designed for regions with higher variability in Table 1.

**Table 1.**
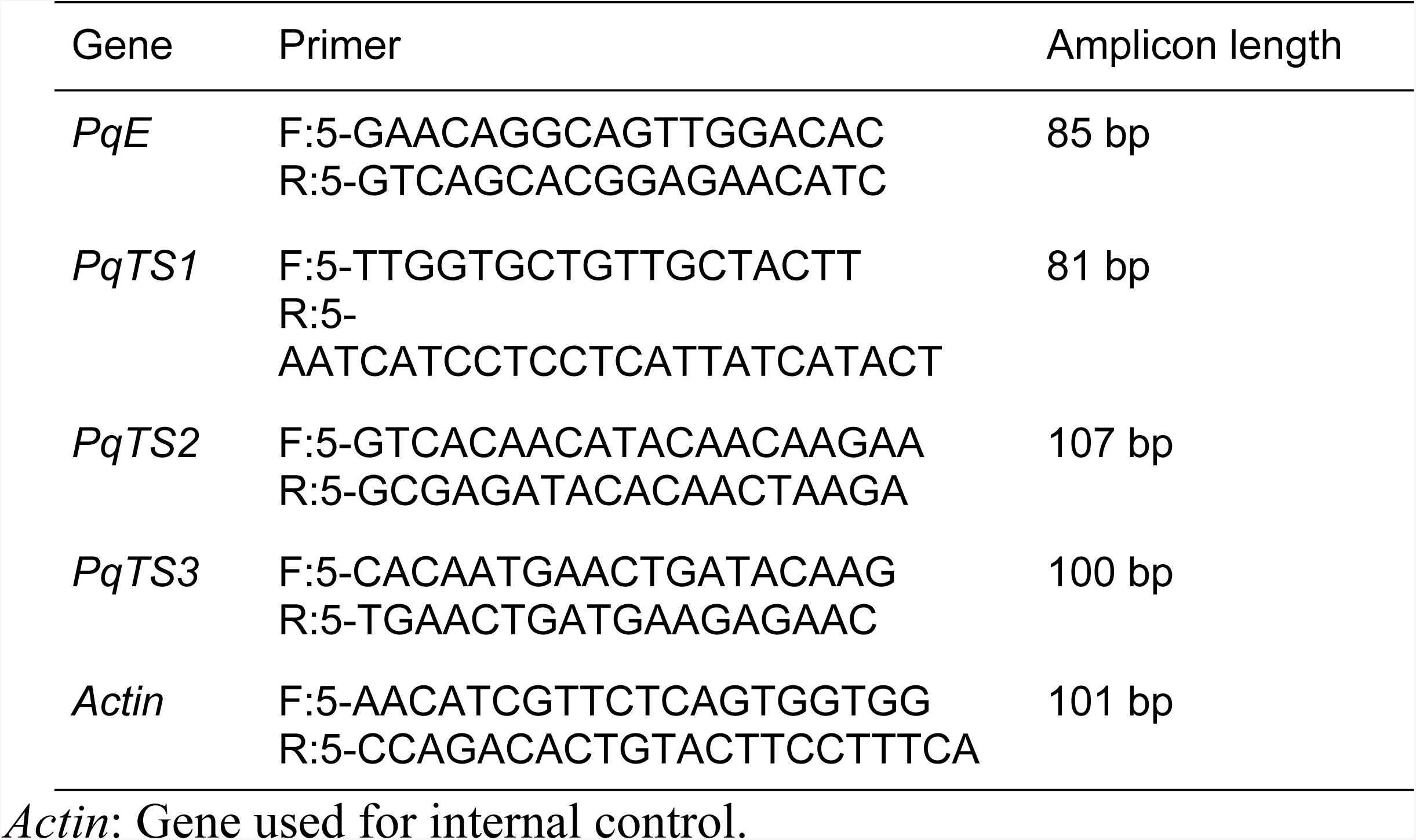
Primers used to amplify four putative genes by real-time PCR.

### HPLC analyses of endogenous polyamines

Endogenous polyamines in the S and R goosegrass biotypes sprayed with paraquat were quantified as described by Flores et al [25]. To extract polyamines, 0.5 g goosegrass leaf tissue was homogenized in 3 ml 5% perchloric acid (PCA) and extracted on ice for 1 h. After centrifugation at 12000 *g* for 20 min at 4°C, the supernatant was collected for analysis using a Varian HPLC system (HPLC; LC-20A, Prominence Series, Shimadzu, Kyoto, Japan). The system comprised a Prostar 210 solvent delivery module, a Prostar 325 UV-Vis Detector, and a 20 μL sample loop (Rheodyne, Rohnert Park, CA, USA). An analytical standard of paraquat for chromatographic analysis was purchased from Sigma Aldrich (St Louis, MO, USA). For each sample, a 100 μL aliquot was analysed for polyamine content after the addition of an equal volume of 5% (w/v) PCA containing 0.12 mM 1,6-diaminohexane (DAH) as an internal standard. A reference solution containing putrescine, spermidine, and spermine was prepared as described above and analysed to establish retention times and signal intensities for each compound and the internal standard. The measurements of polyamine contents in goosegrass were performed according to Wang et al [26].

### Data analysis

Data were analysed by analysis of variance (ANOVA) at a 95% confidence level using Excel 2010 and SPSS 17.0 (SPSS Inc., Chicago, IL, USA). When the ANOVA indicated significant differences between treatments, means were separated using Duncan’s test at P = 0.05.

## Results

### Characterization of four putative genes encoding polyamine transporters

In this study, the characterization of *PqE, PqTS1, PqTS2*, and *PqTS3* was performed by aligning homogeneous sequences from the NCBI website. *PqE, PqTS1, PqTS2,* and *PqTS3* shared high homology (^3^86%) with the *KCS, SYP121, ABCB,* and *Ca*^*2+*^*-ATP* found in foxtail millet (*Setaria italica* (L.) P. Beauv.), respectively, as shown in Table 2. However, the fold change and reads per kilobase of exon model per million mapped reads (PRKM) indicated that *PqE, PqTS1, PqTS2,* and *PqTS3* were more closely related to the paraquat resistance in goosegrass.

**Table 2.** Biological analysis of four putative genes*.

| Gene | Homologous gene | Accession | Description | Query cover | E value | Identity |
| --- | --- | --- | --- | --- | --- | --- |
| *PqE* | *KCS* | XM_012848229.1 | 3-ketoacyl-CoA synthase 11-like | 99% | 0.0 | 88% |
| *PqTS1* | *SYP121* | XM_004981451.4 | syntaxin-121 | 87% | 0.0 | 88% |
| *PqTS2* | *ABCB* | XM_004969589.2 | ABC transporter B family member 4-like | 90% | 0.0 | 86% |
| *PqTS3* | *Ca<sup>2+</sup>-ATP* | XM_004985233.3 | calcium-transporting ATPase 1 or plasma membrane-type | 81% | 0.0 | 87% |

\*More information on the *KCS*, *SYP121*, *ABCB*, and *Ca<sup>2+</sup>-ATP* genes can be found on the NCBI website (http://blast.ncbi.nlm.nih.gov); it should be noted that their sequences were used as query sequences for BLASTN searches.

### Transcript levels of putative polyamine transporter genes after paraquat treatment

The S and R biotypes of goosegrass were treated with paraquat, and the steady-state transcript levels of the selected genes were then determined by RT-PCR. All four putative genes showed different transcription patterns between the S and R biotypes following paraquat treatment (Fig 1A), and the amplification products were of the expected size (Fig 1B).

**Fig 1.**
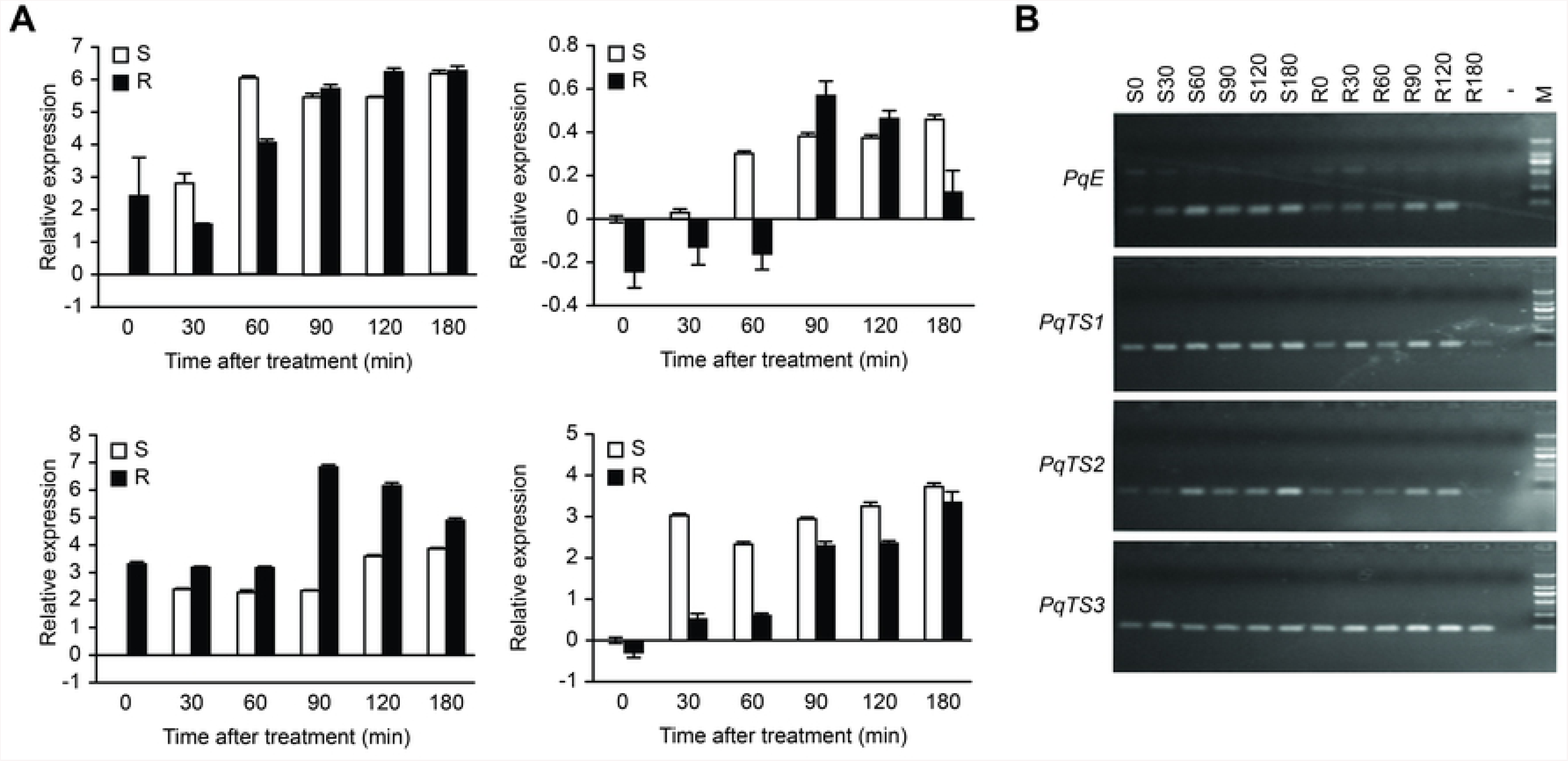
Relative transcript levels of putative genes after paraquat treatment. **A.** Gene transcript levels in susceptible (S) and resistant (R) goosegrass at the indicated times after paraquat treatment. Transcripts were detected by real-time PCR using total RNA extracted from 10-d-old seedlings sprayed with 10/HM paraquat. Error bars represent the SD (n = 3). **B**. Amplification products of 4 putative genes from goosegrass by RT-PCR. For the M:DL 2000 marker, “-” indicates the negative control. Lanes S0, S30, S60, S90, S120, S180, R0, R30, R60, R90, R120 and R180 are samples. R0, R30, R60, R90, R120, and R180 indicate samples of R goosegrass seedlings taken 0, 30, 60, 90, 120 and 180 min after paraquat treatment, while S0, S30, S60, S90, S120 and S180 indicate samples of S goosegrass seedlings taken 0, 30, 60, 90, 120 and 180 min after paraquat treatment, respectively.

The S biotype of goosegrass showed an increase in *PqE* transcript levels 0 min to 90 min after paraquat treatment, but the R biotype of goosegrass did not show an increase in *PqE* transcript levels until 60 min after paraquat treatment. At 0 min, the transcript level of *PqE* in the R biotype was 2.4 times higher than that in the S biotype. Thirty minutes after treatment, *PqE* was significantly upregulated in the S biotype but downregulated to the minimum relative expression in the R biotype. Sixty minutes after treatment, the *PqE* transcript level was remarkably increased in the S biotype (6-fold higher than that in the control), whereas it increased to a lesser extent in the R biotype (4-fold higher than that in the control). Ninety minutes after treatment, the transcript level of *PqE* did not differ significantly between the S and R biotypes (5.54-fold and 5.73-fold higher than that in the control, respectively). Finally, stable expression of *PqE* was detected in both ecotypes 120 min and 180 min after paraquat treatment.

The transcript level of *PqTS1* was higher in the R biotype than in the S biotype at each stage 0-180 min after paraquat treatment. This level remained steady in the R biotype until 60 min after paraquat treatment, peaked at 90 min, and then decreased at 120 and 180 min. In the S biotype, the transcript level of *PqTS1* remarkably increased at 30 min; thereafter, it remained stable until 90 min and then increased slightly until 180 min after the paraquat treatment.

The *PqTS2* transcript level showed detectable changes in both the R and S biotypes. The *PqTS2* transcript level was approximately 0.24-fold lower in the R biotype than in the S biotype at 0 min without paraquat treatment. The *PqTS2* transcript levels were upregulated in the S biotype and downregulated in the R biotype 60 min after paraquat treatment. However, 90 and 120 min after paraquat treatment, the *PqTS2* transcript level was higher in the R biotype than in the S biotype.

The paraquat treatment increased *PqTS3* in both biotypes. At 0-90 min after paraquat treatment, *PqTS3* was significantly higher in the S biotype than in the R biotype. Although the transcript level of *PqTS3* in the R biotype was 0.32-fold lower than that in the S biotype at 0 min, it steadily increased in the R biotype after 30 to 180 min. In the S biotype, *PqTS3* transcript levels remained relatively steady 30 to 180 min after paraquat treatment.

### Endogenous polyamine contents in goosegrass biotypes

We used HPLC to detect differences in endogenous polyamine contents between S and R seedlings 0 to 180 min after paraquat treatment (S1 Fig). Polyamine levels in S goosegrass and R goosegrass after a paraquat treatment are shown (Fig 2). In the S biotype, the putrescine content slightly fluctuated 0 and 90 min after paraquat treatment and then significantly increased after 120 min. The spermidine and spermine contents in S seedlings did not show significant differences from the control at any time after paraquat treatment. In contrast, there were significant changes in polyamine contents, especially spermidine content, in R seedlings after paraquat treatment. After paraquat treatment, the spermidine concentration in R seedlings increased 421.04-fold at 30 min and 346.14-fold at 60 min, but at 90 min, it decreased to an infinitesimal level. In the R biotype, putrescine levels increased 30 min after paraquat treatment and remained steady until 180 min. The spermine content significantly increased in R seedlings 0 to 90 min after paraquat treatment and then decreased 90 to 180 min after treatment. Overall, the endogenous polyamine content in the R biotype was higher than that in the S biotype after paraquat treatment in S1 Table.

**Fig 2.**
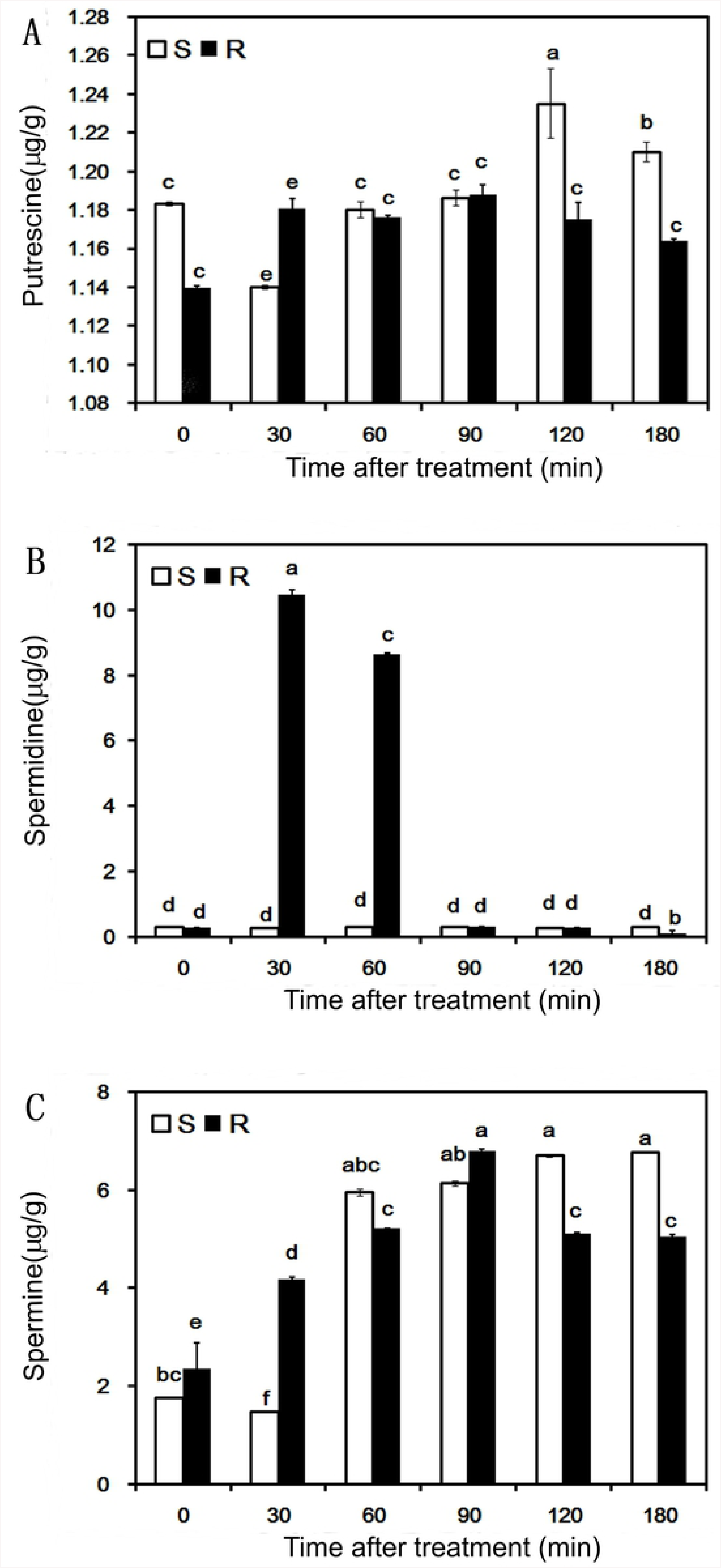
Polyamine levels in susceptible (S) and resistant (R) goosegrass after a paraquat treatment. Putrescine (A), spermidine (B), and spermine (C) were extracted from shoots of R goosegrass (black bar) and S goosegrass (white bar) seedlings 0, 30, 60, 90, 120, and 180 min after spraying with paraquat and then quantified by HPLC. Different lowercase letters indicate significant differences at P<0.05 (t-test).

## Discussion

Paraquat and polyamines show similar uptake characteristics in animal and plant systems, suggesting that paraquat uptake is mediated by polyamine transporters [27]. Paraquat toxicity in plants may result when paraquat is taken up by cells via a polyamine transporter with a function under stringent negative regulation by spermidine [28]. Moreover, Kurepa et al. reported that spermidine was the most effective polyamine in protecting *Arabidopsis* against paraquat toxicity, possibly because paraquat is taken up by cells via a polyamine transporter with a function negatively regulated by spermidine [28]. This speculation is based on the fact that paraquat exists as a divalent cation that is structurally similar to *α,ω-*diamines and is taken up by cells via polyamine transporters in aqueous solution.

*PqE* shares high homology with the *β-ketoacyl CoA synthase* (*KCS*) gene, which catalyses an essential reaction in the fatty acid elongation process. *PqTS1* shares high homology with *SYP121,* which encodes a protein involved in the transport of secretory vesicles at the plasma membrane [29]. Transgenic plants overexpressing *MerC-SYP121* in the plasma membrane were more resistant to cadmium than was the wild type [30]. Moreover, *SYP121/PEN1* mutants showed enhanced disease susceptibility in compatible host-pathogen interactions [31]. We found that *PqTS2* shares high homology with *ABCB*, which encodes a member of the ABC transporter B family (ATP-binding cassette) of proteins. This finding suggests that *PqTS2* has functions similar to those of the homogenous *MDR1*/*ABCB1* gene, which is also function at an efflux transporter of paraquat in protection against subsequent toxicity in humans and mice [32]. *PqTS3* shares high homology with Ca^2+^-ATP, which encodes ATPase 1 or plasma membrane-type Ca^2+^-ATP. Paraquat inhibited Ca^2+^ influx across the plasmalemma, and the phenylacetic acid transport system responsible for the movement of paraquat across the plasmalemma was relatively specific to divalent cationic molecules with a distinct charge distribution [33].

Our research corroborates the results of previous studies showing that paraquat treatment can decrease the relative expression level of *PqE.* The *PqE* transcript levels decreased while the spermidine and spermine contents significantly increased in the R biotype after paraquat treatment (Figs1and 2). This result may be because paraquat was bound to the polyamine transporter, thereby hindering polyamine transport. The resulting accumulation of polyamines could then reduce the activities of polyamine synthases. Thus, *PqE* might be involved in both polyamine and paraquat resistance. Secondly, *PqTS1* showed higher transcript levels and more sustained expression in the R biotype than in the S biotype after the paraquat treatment. Increases in spermidine and spermine contents alleviated the toxic effects of paraquat in R goosegrass 0 to 60 min after spraying, suggesting that *PqTS1* played a role in vesicular membrane transport of spermidine and spermine in cells. Thus, the upregulation of *PqTS1* expression was an important part of the resistance response to paraquat in goosegrass. However, further studies on *PqTS1* are required to clarify its role in paraquat resistance. Thirdly, the transcript level of *PqTS2* in the R biotype decreased 0 to 60 min after the paraquat treatment, and the subsequent detoxification effect of spermidine decreased 90–120 min after spraying with paraquat, possibly indicating that the increased expression of *PqTS2* 90 min after the paraquat treatment affected paraquat transport from goosegrass leaves. The concentration of putrescine did not differ significantly between the R and S biotypes 0–60 min after the paraquat treatment. This result was consistent with the finding that paraquat completely inhibited putrescine uptake but not spermidine uptake by the spermidine-preferential ABC transporter in *E. coli* [34]. The spermidine content was closely related to the *PqTS2* transcript levels in goosegrass in our study, suggesting that *PqTS2* plays a key role in the molecular mechanism of paraquat resistance. However, further research is required to explore the function of Spm in paraquat resistance. Finally, the relative expression of *PqTS3* was significantly different between the S biotype and R biotype after paraquat treatment in our study. Based on this finding, we speculate that the putative gene *PqTS3* takes part in the regulation of the levels of spermidine and spermine to alleviate paraquat toxicity in R goosegrass, which could lead to later signal reception of paraquat in R goosegrass than in S goosegrass.

## Conclusion

Endogenous polyamines actually play a vital role in paraquat resistant in goosegrass, especially the spermidine. Meanwhile, 4 genes have been identified in this paraquat-resistance weed, which encode putative components of polyamine transport and synthesis systems.

## Acknowledgements

This work was supported by the National Natural Science Foundation of China (Nos. 31471788, 31272054). We are grateful to the National Natural Science Foundation of China for supporting this study.

## Author Contributions

Y.C. conceived and designed the experiments. Q.L. wrote the manuscript. Q.L. and J.W. performed most of the experiments. Z.D. and X.S. contributed reagents and materials. All authors read and approved the final manuscript.

## Supporting information

**S1 Table.** Paraquat effects on the endogenous polyamine contents in goosegrass.

| Type | Polyamine content ($\mu\text{g g}^{-1}\text{FW}^{-1}$) | | | | | |
| --- | --- | --- | --- | --- | --- | --- |
|  | 0 min | 30 min | 60 min | 90 min | 120 min | 180 min |
| Susceptible (S) goosegrass | 32.48i | 28.84j | 74.08e | 76.05d | 82.11c | 82.81c |
| Resistant (R) goosegrass | 37.79h | 158.24a | 150.36b | 83.04c | 65.73f | 63.24g |

Polyamines: Putrescine, spermidine and spermine. Different lowercase letters indicate significant differences at $P < 0.05$ (t-test).

**S1 Fig. HPLC wave peak diagram of polyamine contents in susceptible (S) and resistant (R) goosegrass after paraquat treatment**.

R0, R30, R60, R90, R120, and R180 indicate samples of the resistant goosegrass seedlings taken 0, 30, 60, 90, 120 and 180 min after paraquat treatment, while S0, S30, S60, S90, S120 and S180 indicate samples of susceptible goosegrass seedlings taken 0, 30, 60, 90, 120 and 180 min after paraquat treatment, respectively. Put represents putrescine, Spd represents spermidine, and Spm represents spermine.

## References

1. Jóri B, Soós V, Szegő D, Páldi E, Szigeti Z, Rácz I, et al. Role of transporters in paraquat resistance of horseweed *Conyza canadensis* (*L.*) cronq. Pestic Biochem Physiol. 2007;88: 57–65.

2. Dong S, Hu H, Wang Y, Xu Z, Zha Y, Cai X, et al. A pqr2 mutant encodes a defective polyamine transporter and is negatively affected by ABA for paraquat resistance in Arabidopsis thaliana. J Plant Res. 2016;129: 899–907.

3. Xi J, Xu P and Xiang CB. Loss of AtPDR11, a plasma membrane-localized ABC transporter, confers paraquat tolerance in Arabidopsis thaliana. Plant J. 2012; 69: 782–791.

4. Tseng T, Ou J and Wang C. Role of the ascorbate–glutathione cycle in paraquat tolerance of rice. Weed Sci. 2013;61: 361–373.

5. Waris G and Ahsan H. Reactive oxygen species: role in the development of cancer and various chronic conditions. J Carcinog. 2006;5: 14–14.

6. Tsuji K, Hosokawa M, Morita S, Miura R, Tominaga T and Kudsk P. Resistance to paraquat in Mazus pumilus. Weed Res. 2013; 53: 176–182.

7. Powles SB and Yu Q. Evolution in action: plants resistant to herbicides. Annu Rev Plant Biol. 2010; 61:317–347.

8. Hawkes R. Mechanisms of resistance to paraquat in plants. Pestic Manage Sci. 2014;70: 1316–1323.

9. Pandolfi C, Pottosin I, Cuin T, Mancuso S and Shabala S. Specificity of polyamine effects on NaCl-induced ion flux kinetics and salt stress amelioration in plants. Plant Cell Physiol. 2010; 51: 422–434.

10. Hart JJ, Ditomaso JM, Linscott DL and Kochian LV. Transport interactions between Paraquat and polyamines in roots of intact maize seedlings. Plant Physiol. 1992;99: 1400–1405.

11. Benavides M, Gallego S, Comba M and Tomaro M. Relationship between polyamines and paraquat toxicity in sunflower leaf discs. Plant Growth Regul 31:215-224 (2000).

12. Chang C and Kao C. Paraquat toxicity is reduced by polyamines in rice leaves. Plant Growth Regul 22:163-168 (1997).

13. Soar CJ, Preston C, Karotam J and Powles SB. Polyamines can inhibit paraquat toxicity and translocation in the broadleaf weed Arctotheca calendula. Pestic Biochem Physiol. 2004;80: 94–105.

14. Dinis-Oliveira RJ, Duarte JA, Sanchez-Navarro A, Remiao F, Bastos ML and Carvalho F. Paraquat poisonings: mechanisms of lung toxicity, clinical features, and treatment. Crit Rev Toxicol. 2008;38: 13–71.

15. Szigeti Z. Mechanism of paraquat resistance-from the antioxidant enzymes to the transporters. Acta Biol Szeged. 2005;49: 177–179.

16. Jung HI, Kuk YI, Kim HY, Back K, Lee DJ, Lee S, et al. Resistance levels and fitness of protoporphyrinogen oxidase (PROTOX) inhibitor-resistant transgenic rice in paddy fields. Field Crops Res. 2010;115: 125–131.

17. Li J, Mu J, Bai J, Fu F, Zou T, An F, et al, Yang S and Zuo J. Paraquat resistant1, a golgi-localized putative transporter protein, is involved in intracellular transport of paraquat. Plant Physiol. 2013;162: 470–483.

18. Zhang C, Feng L and Tian XS. Alterations in the 5’ untranslated region of the 5- enolpyruvylshikimate-3-phosphate synthase (EPSPS) gene influence EPSPS overexpression in glyphosate-resistant *Eleusine indica*. Pestic Manage Sci. 2018;74: 2561–2568.

19. Lii R, Steed S and Stall W. Confirmation and control of a paraquat-tolerant goosegrass (Eleusine indica) biotype. Weed Technol. 2002;16: 309–313.

20. Seng C, Lun L, Chathye S and Sahid I. Initial report of glufosinate and paraquat multiple resistance that evolved in a biotype of goosegrass (Eleusine indica) in Malaysia. Weed Biol Manag. 2010;10: 229–233.

21. An J, Shen X, Ma Q, Yang C, Liu S and Chen Y. Transcriptome profiling to discover putative genes associated with paraquat resistance in goosegrass (Eleusine indica L.). PLoS One. 2014; 9: e99940.

22. Shen X, Hu F, Chen Y, Li Y and Han C. Preliminary study on resistance level of Eleusine indica to paraquat. Southwest China Journal of Agricultural Sciences. 2016;29: 1875–1878.

23. Altschul SF, Gish W, Miller W, Myers EW and Lipman DJ. Basic local alignment search tool. J Mol Biol. 1990;215: 403–410.

24. Thompson JD, Higgins DG, Gibson TJ and Clustal W. Improving the sensitivity of progressive multiple sequence alignment through sequence weighting, position-specific gap penalties and weight matrix choice. Nucleic Acids Res. 1994;22: 4673–4680.

25. Flores HE and Galston AW. Analysis of polyamines in higher plants by high performance liquid chromatography. Plant Physiol. 1982;69: 701–706.

26. Wang F and Xue Y. Measurement of polyamine contents in plant tissue. Plant Physiol Commun 1988;1: 39–41.

27. Fujita M, Fujita Y, Iuchi S, Yamada K, Kobayashi Y, Urano K, et al. Natural variation in a polyamine transporter determines paraquat tolerance in *Arabidopsis*. Proc Natl Acad Sci USA 2012;109: 6343–6347.

28. Kurepa J, Smalle J, Van Montagu M and Inze D. Polyamines and paraquat toxicity in *Arabidopsis thaliana*. Plant Cell Physiol. 1998;39: 987–992.

29. Kiyono M, Oka Y, Sone Y, Nakamura R, Sato MH, Sakabe K, et al. Bacterial heavy metal transporter MERC increases mercury accumulation in *Arabidopsis thaliana*. Biochem Eng J. 2013;71: 19–24.

30. Kiyono M, Oka Y, Sone Y, Tanaka M, Nakamura R, Sato MH, Pan-Hou H, Sakabe K and Inoue K. Expression of the bacterial heavy metal transporter MERC fused with a plant SNARE, SYP121, in *Arabidopsis thaliana* increases cadmium accumulation and tolerance. Planta. 2012; 235: 841–850.

31. Collins NC, Thordal-Christensen H, Lipka V, Bau S, Kombrink E, Qiu JL, Huckelhoven R, Stein M, Freialdenhoven A, Somerville SC and Schulze-Lefert P. SNARE-protein-mediated disease resistance at the plant cell wall. Nature. 2003;425: 973–977.

32. Wen X, Gibson CJ, Yang I, Buckley B, Goedken MJ, Richardson JR and Aleksunes LM. MDR1 transporter protects against paraquat-induced toxicity in human and mouse proximal tubule cells. Toxicol Sci. 2014;141:475–483.

33. Hart JJ, Ditomaso JM, Linscott DL and Kochian LV. Investigations into the cation specificity and metabolic requirements for paraquat transport in roots of intact maize seedlings. Pestic Biochem Physiol. 1993;45: 62–71.

34. Kashiwagi K, Hosokawa N, Furuchi T, Kobayashi H, Sasakawa C, Yoshikawa M and Igarashi K. Isolation of polyamine transport-deficient mutants of *Escherichia coli* and cloning of the genes for polyamine transport proteins. J Biol Chem. 1990;265: 20893–20897.

